# A statistical approach reveals designs for the most robust stochastic gene oscillators

**DOI:** 10.1101/025056

**Authors:** Mae Woods, Miriam Leon, Ruben Perez-Carrasco, Chris P. Barnes

**Affiliations:** Department of Cell and Developmental Biology, University College London; Department of Mathematics, University College London; Department of Genetics, Evolution and Environment, University College London

## Abstract

The engineering of transcriptional networks presents many challenges due to the inherent uncertainty in the system structure, changing cellular context and stochasticity in the governing dynamics. One approach to address these problems is to design and build systems that can function across a range of conditions; that is they are robust to uncertainty in their constituent components. Here we examine the parametric robustness landscape of transcriptional oscillators, which underlie many important processes such as circadian rhythms and the cell cycle, plus also serve as a model for the engineering of complex and emergent phenomena. The central questions that we address are: Can we build genetic oscillators that are more robust than those already constructed? Can we make genetic oscillators arbitrarily robust? These questions are technically challenging due to the large model and parameter spaces that must be efficiently explored. Here we use a measure of robustness that coincides with the Bayesian model evidence combined with an efficient Monte Carlo method to traverse model space and concentrate on regions of high robustness, which enables the accurate evaluation of the relative robustness of gene network models governed by stochastic dynamics. We report the most robust two and three gene oscillator systems, plus examine how the number of interactions, the presence of auto-regulation, and degradation of mRNA and protein affects the frequency, amplitude and robustness of transcriptional oscillators. We also find that there is a limit to parametric robustness, beyond which there is nothing to be gained by adding additional feedback. Importantly, we provide predictions on new oscillator systems that can be constructed to verify the theory and advance design and modelling approaches to systems and synthetic biology.

## Introduction

A major challenge facing the progress of synthetic biology is the design and implementation of systems that function in the face of fluctuating cellular environments. While it is widely accepted within the field that the task of constructing and rewiring pathways is tractable, predicting *in silico* how an implemented system will behave *in vivo* under different cellular conditions remains a huge challenge.^1^ Robust systems perform their function over a wide range of parameters and external influences. If we could design and build synthetic systems that are robust, then not only would the systems have a higher probability of functioning, but we would also enhance their predictability. Robustness in the context of biological systems has been intensively studied for almost two decades.^2–6^ Approaches to studying this in biological systems often utilise the frameworks of feedback and robust control.^7^ It is well known that feedback mechanisms can increase the robustness of a biological system^8,9^ and there are tradeoffs between robustness and performance, fragility and efficiency.^10,11^ Although some biological systems have been shown to be structurally robust - that is the underlying biochemical rate parameters have little effect on the system stability properties^12^ - in general we expect system behaviour to depend heavily on the biochemical parameters.^13^

Biological oscillators have been studied extensively as they form the core of many crucial biological processes such as circadian rhythms and the cell cycle. Oscillating systems also serve as a model for the understanding and engineering of complex and emergent phenomena. Various synthetic systems have been implemented both *in vivo* and *in vitro*.^14–20^ More complex behaviours have been constructed, enabling synchronisation over multiple scales, and entrainment by external signals.^21–23^ There has been much theoretical study of biological oscillators (for reviews see refs^24–26^). Specific work has been done on noise attenuation,^27^ motifs capable of oscillation,^28–31^ robustness^32–34^ and the role of positive and negative feedback.^35,36^ Feedback in natural circadian oscillators has also been studied.^37,38^

Despite this body of work, a comprehensive study of the robustness of transcriptional oscillators has not been performed because of the technical challenges it poses. Traditional mathematical approaches can elucidate general design principles and are of great importance^25^ but these techniques generally simplify systems down to a handful of parameters. More contemporary methods can also explore model and phenotype space and are less restricted in model size^31,39^, but they rely on the analysis of deterministic dynamics and cannot handle the full complexity of realistic stochastic biological systems. Therefore, to develop more predictable design and modelling frameworks that can calculate realistic estimates of system properties - including robustness - requires approaches that can handle a large number of parameters, parametric uncertainty and stochastic dynamics. This can be achieved using sequential Monte Carlo methods.^40,41^ Here we extend the Monte Carlo framework to include model space exploration. The novelty in our approach is that the algorithm spends time in models and parameters in direct proportion to their robustness, and thus focuses in on interesting regions of joint model-parameter space. This avoidance of enumeration of all possibilities allows us to address more interesting questions and to assess robustness in a quantitative manner.

We apply this novel framework to investigate the robustness of transcriptional oscillators, an outline of which is given in Figure 1. We examine two main questions regarding the robustness of stochastic transcriptional oscillators: Can we build genetic oscillators that are more robust than those already constructed? Can we make genetic oscillators arbitrarily robust? We find that the most robust two gene oscillators that can provide regular oscillations are of a type already constructed.^17^ We also examine the ring oscillator - the repressilator being the classic synthetic implementation^14^ - and find that different activation reactions, in addition to positive auto-regulation,^35^ can increase its robustness. We also determine the topologies that give rise to the most robust three gene systems and find that in general they are more robust than the simple two gene and ring oscillators. The frequency, amplitude and robustness of all transcriptional oscillators, independent of topology, depends strongly on the rates of degradation of the species involved. Finally we find that the number of regulatory interactions increases oscillator robustness up to a plateau, beyond which there is no increase in robustness, which has wide implications for the construction of complex synthetic systems.

**Figure 1:**
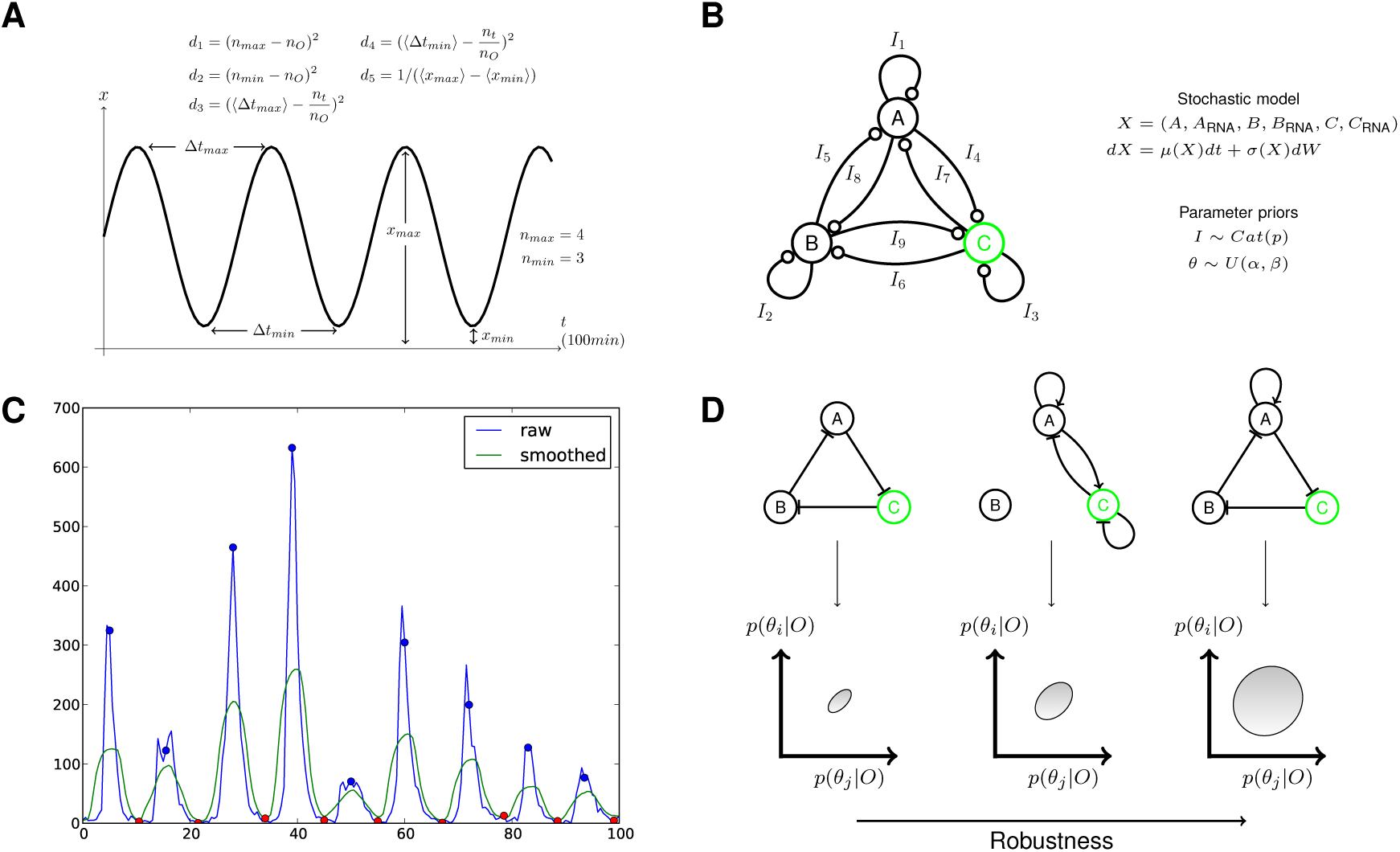
Outline of the method. (A) The objective behaviour is specified through a set of summary statistics and distances on the summaries (see Materials and Methods for a description of the terms). (B) Model space is defined through a fully connected network. A mapping from the graphical network to a stochastic model is defined together with a prior on the parameters and priors on the allowed networks. (C) Simple signal processing methods are used to extract features from model simulations. The blue and red circles indicate identified maxima and minima respectively. (D) As the algorithm proceeds the (multi-dimensional) objective is achieved via small increments using sequential Monte Carlo. The final output of the algorithm is a set of models that satisfy the objective and maximise robustness.

## Results and discussion

### The most robust two-node oscillators combine positive and negative auto-regulation

We searched model space for the most robust two-node oscillators and considered all possible regulatory interactions resulting in 3^4^ = 81 different models (Figure 2A). We found that only five models were capable of oscillations at the specified frequency (Materials and Methods), which we denote by M1-5 (Figure 2B). M1 and M2, which are mirror images, account for around 93% of the posterior model space and have the structure of an amplified negative feedback loop with negative auto-regulation on the repressor and positive auto-regulation on the activator. The other three systems all contain negative auto-regulation of protein A (the protein that does not serve as the output), a well known oscillatory motif, but show high levels of stochasticity. Interestingly the topology represented by M1 and M2 formed the basis for the robust oscillator constructed by Stricker *et. al*.^17,42^ We also note the absence of the delayed negative feedback oscillator with no negative auto-regulation on the repressor, which is consistent with the observation that it cannot produce sustained oscillations.^15,20,43^ The last oscillator system, denoted here by M5, has also been constructed and shown to produce more stochastic behaviour than the amplified negative feedback topology of M1 and M2.^17^ These findings demonstrate that the modelling framework can reconstitute empirical findings in real synthetic oscillators.

**Figure 2:**
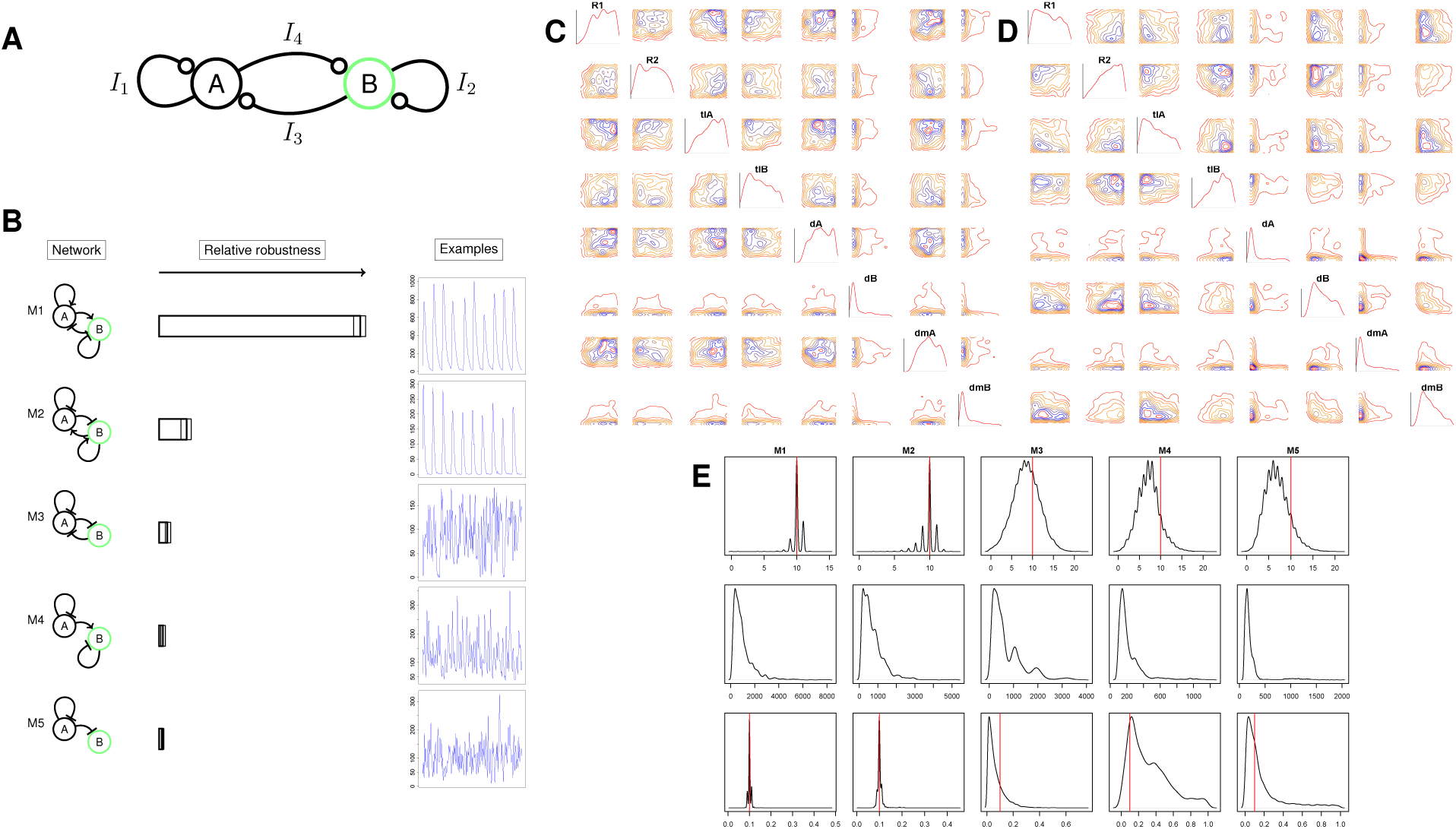
The most robust two-gene oscillators. (A) The regulatory interactions considered for the systems. The output protein, where oscillations are required, is shown in green. (B) The most robust two-node oscillators. The error bars indicate the variability in the marginal model posteriors over three separate runs (minimum, median, maximum). On the right are representative example time series of the time course behaviour.. (C,D) Posterior parameter distributions for the top two models, M1 and M2, for production, translation and degradation rates of proteins *A* and *B* (*R*_1_, *R*_2_, *tl*_A_, *tl_B_, d_A_,d_B_*) and the degradation rates of *A_mRNA_* and *B_mRNA_* (*d_mA_, d*_mB_). The posteriors are plotted on their prior range of (0,10000) for the production rates and (0,10) for the decay rates. (E) Model checking of the stochastic systems by resampling and re-simulation under the posterior distribution. The top, middle and bottom plots correspond to the number of oscillations, the amplitude and the maximal frequency of the Fourier spectrum respectively.

Since our approach examines the dynamics of a specified protein (in this case B), we can elucidate how targeting different output nodes affects robustness. In the two-node oscillator we see that system M1 is around 8 times more robust than system M2 (Bayes factor ≈ 8) despite the fact that the only difference is the output node. This can be understood in terms of the dynamics of the relaxation oscillator. The repressor generally has slower dynamics and reaches higher numbers of mRNA and protein molecules in comparison to the activator (Supplementary Figure 4). Placing the output on the activator (M2) and requiring a minimum amplitude on the resultant oscillations forces the levels of repressor to reach a higher amplitude than is necessary when the output is placed on the repressor (M1). This can also be seen by examining the system size (total number of mRNA and protein molecules) which is larger in M2 (Supplementary Figure 4). These additional requirements on the dynamics constrains the parameters of M2. The one and two dimensional marginal posteriors for a subset of the parameters are given for the two oscillator systems in Figure 2C-D. The parameter posterior distributions for M1 are closer to their prior distributions, with *tl_A_, tl_B_, d_A_* and *d_mA_* appearing to be virtually unconstrained. These flat distributions indicate that a larger fraction of the parameter space in M1 can give rise to oscillations when compared to M2. We also note that M2 requires a higher promoter strength (*R*_2_) upstream of gene B. These posterior distributions can also be used to aid the design process by providing information on the required biochemical properties to create functional and robust systems.

Defining oscillations in stochastic systems is non-trivial,^17^ and separating true oscillators from excitable systems can be difficult. To investigate the stochasticity of the five systems, and whether they are truly oscillators, we applied a statistical model checking procedure, whereby the posterior parameter distributions are resampled and the systems re-simulated to examine the resultant performance. Figure 2E shows the results of the resampling and recalculation of the number of oscillations (top) and amplitude (middle). M1 and M2 have a large peak at ten pulses in the 100 minute time period, showing a consistent and reliable performance. In contrast M3–5 show much wider distributions indicating that they are more stochastic. There are also differences between the oscillator amplitude properties with M4 and M5 showing lower amplitude oscillations. To verify the frequency properties in an unbiased manner we also calculated the maximal frequency of the Fourier spectrum which was not included in the objective summary statistics (Figure 2E bottom). Taken together, these results suggest that M1 and M2 have very reproducible frequency properties, both M4 and M5 are true oscillators albeit with high stochasticity and low amplitude, and M3 is most likely to be an excitable system. From an engineering point of view, we are only interested in regular oscillators, and so can exclude M3-M5 as failing our design objectives.

To examine how the robustness changes under specification of the objective behaviour, we compared the two-node systems under the objectives S1 (fixed frequency) and S2 (variable frequency, regular oscillations) (Supplementary Figure 5). We found almost identical results indicating that in this particular system, and under our prior assumptions, the difference between these objectives is minimal.

### Different activation reactions increase the robustness of the ring oscillator

Next we examined how one can increase the robustness of the three gene ring oscillator by the incorporation of additional feedback interactions. The setup for the model is shown in Figure 3A. The three negative feedback interactions of the standard ring oscillator were fixed, with additional activating and repressing interactions allowed, giving a model space of 3^6^ = 729 models. The distribution of the Bayes factor with respect to the standard ring oscillator for the 30 most robust topologies is shown in Figure 3C. The twelve most robust networks are shown in Figure 3D. Additional positive auto-regulation clearly increases the robustness of the ring oscillator by as much as an order of magnitude (Bayes factor ≈ 10 in Figure 3C). This is in agreement with previous work that showed that a ring oscillator with a single positive auto regulatory feedback loop could achieve more robust oscillatory behaviour than the ring oscillator with or without a single negative auto-regulatory feedback loop.^35^ The benefit of this framework is that we can quantitatively estimate the gain in modifying the original design and judge its worth; a Bayes factor of > 10 indicates strong evidence. We also find that including an additional activation from gene *A* to *C* can significantly increase robustness. This can be seen more clearly by adopting the notion of inclusion probabilities, which rank the regulatory interactions by their probability of occurring in the ensemble of systems in the posterior distribution (Figure 3E). Here we clearly see that including auto-regulation on gene *A* has the best chance of increasing robustness, followed by activation from gene *A* to *C*. Interestingly this type of activating interaction within a ring oscillator was found in the arabidopsis circadian oscillator.^37^

**Figure 3:**
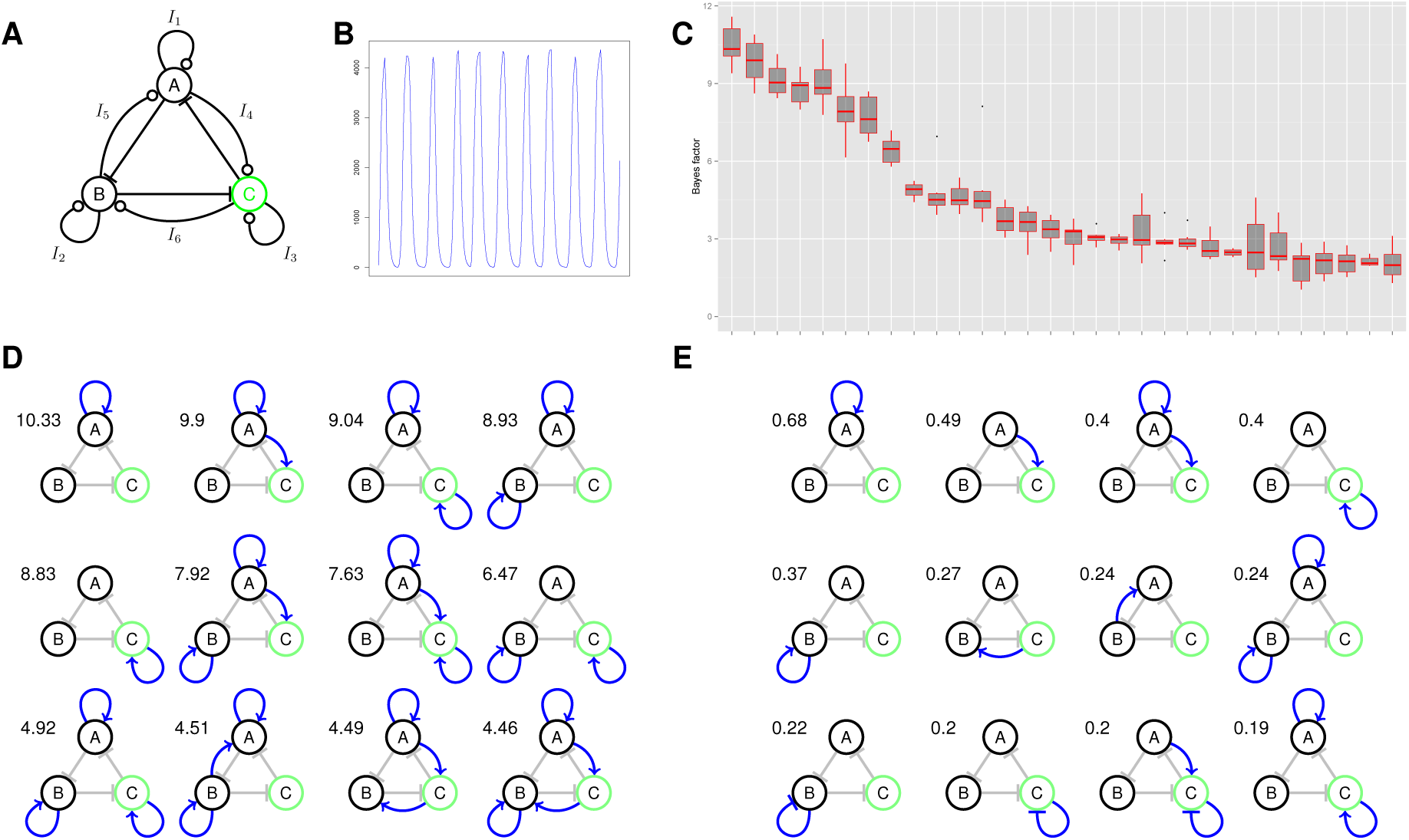
Increasing the robustness of the ring oscillator (A) The regulatory interactions considered for the ring oscillator system. The output protein, where oscillations are required, is shown in green. (B) A representative time series from this system with an oscillation every 10 minutes. (C) The distribution of the 30 most robust topologies. In the box plot, the central bar indicates the median estimate and the upper and lower quartiles correspond to the top and bottom of the box. The points correspond to outliers. The y axis represents the Bayes factor with respect to the ring oscillator (here corresponding to the system with no additional feedback interactions.) (D) The most robust twelve oscillators. The number given next to each network gives the median Bayes factor compared to the basic ring oscillator. (E) The top twelve regulatory interactions ranked by inclusion probability.

We created a graph in which nodes represent auto-regulatory motifs and edges connect nodes related by the addition or removal of a single auto-regulatory interaction (Supplementary Figure 6). We find that positive auto-regulation is associated strongly with robustness, with negative auto-regulation associated with the least robust systems. Interestingly the relationship between the addition of positive auto-regulatory feedback and robustness is non-monotonic and the addition of three such interactions appears to decrease robustness significantly. Examination of the posterior parameter distributions for the production and decay rates of the ring oscillator and the ring oscillator with positive auto-regulation on gene A (Supplementary Figure 7), shows that for these systems to function at this oscillator frequency, the values of the decay rates are very important. In the former, the decay rates of protein C and all mRNA species are constrained to be high. In the latter only the decay rates for C (protein and mRNA) are constrained to be high. Interestingly this is in stark contrast to the two-node oscillators which in general need to have low decay rates for the mRNA and protein species. We again examined robustness under the objectives S1 (fixed frequency) and S2 (variable frequency, regular oscillations) and obtained very similar results (Supplementary Figure 8). We also directly examined the correlation between model robustness under the two objectives, which we found to be reasonably high (Pearson correlation 0.76, Supplementary Figure 9), though the agreement increases with robustness. The posterior distributions of the species decay terms show looser requirements on the decay rates of mRNA and protein species of gene C.

A natural question that arises is how does the robustness of the two gene and ring oscillators compare. We addressed this by using a reduced network topology and prior model space (Figure 4). We find that the two gene oscillator is more robust than the basic ring oscillator though only weakly (Bayes factor ≈ 2). However, the addition of the positive feedback auto-regulatory loop clearly out performs both with a Bayes factor of ≈ 10.8 and 5.7 compared to the ring oscillator and two-node oscillator respectively.

**Figure 4:**
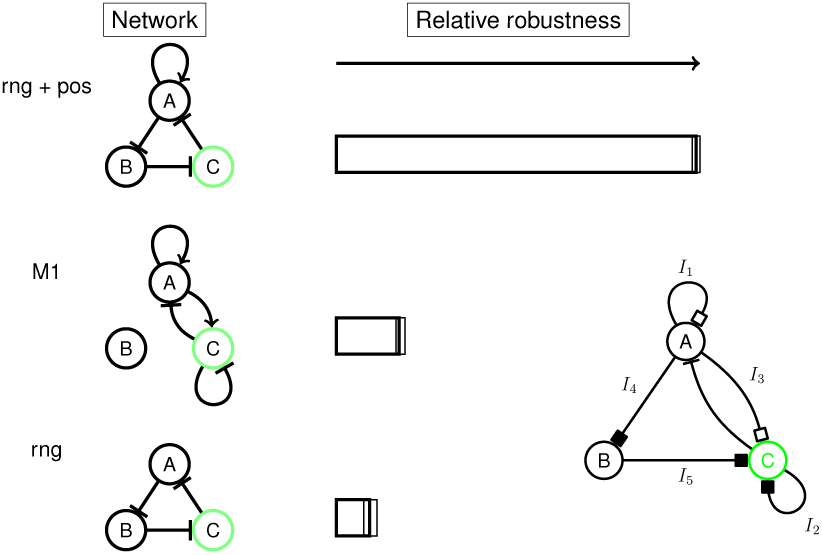
Direct comparison of the relative robustness of the hasty oscillator, the ring oscillator and the ring oscillator with a single positive feedback. Inset: The network used for the direct comparisons. Black squares represent either none or negative regulation ({−1, 0}) and white squares represent either none or positive regulation ({0, +1})

### Robustness of three-node oscillators is achieved through combinations of oscillating motifs

In the previous section we investigated ring oscillators, which are a particular case of the more general three-node oscillator. Here we directly addressed the question of which three gene oscillators give the most robust systems. Rather than considering individual topologies, we explored the more general landscape of possible systems by examining the core architectures. We considered the general three-node network given in Figure 5A with 48 parameters, but restricted the model prior space by setting the prior probability of half the symmetric systems to be zero, resulting in 9963 independent networks (see Supplementary Information). Given the similar results between the fixed frequency objective (S1) and the regular oscillation objective (S2), we used the latter (the phenotypes are shown in the heat map in Figure 5B). The resultant systems were uniquely classified into categories dependent on the core architecture (i.e. ignoring the auto-regulatory interactions). We found in total 43 out of a possible 138 architectures that were reproducible, with robustness spanning over two orders of magnitude. Figure 5C shows the relative robustness of these categories, scaled by the number of topologies in each category. The top ten network topologies are shown in Figure 5D and roughly span an order of magnitude in robustness (Bayes factor ≈ 10).

**Figure 5:**
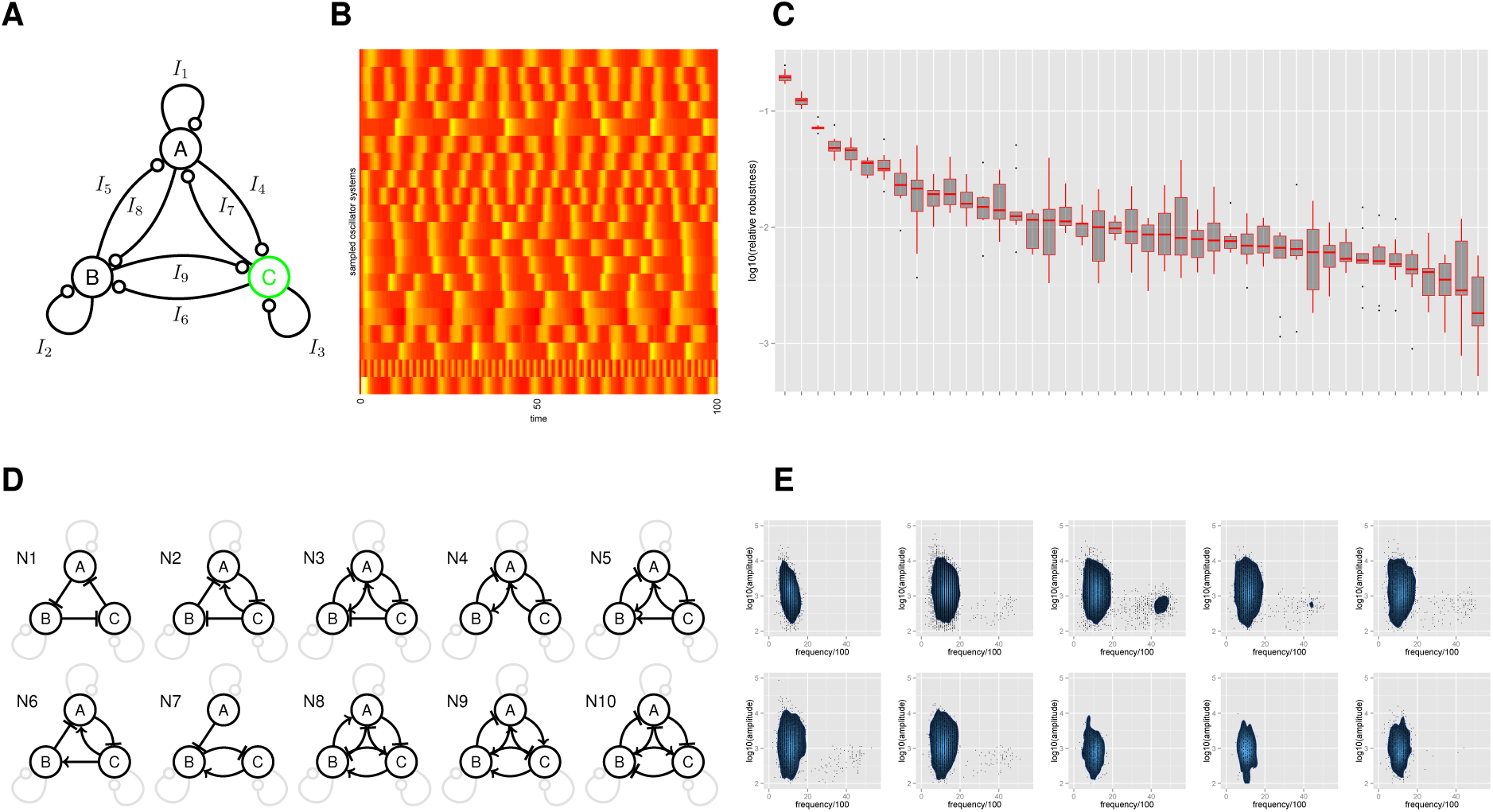
The most robust three-gene networks. (A) The regulatory interactions considered for the three-node oscillator system. The output protein, where oscillations are required, is shown in green. (B) Example phenotypes of the resulting oscillators. Each row of the heat map represents one sampled oscillator system. The yellow and red colours depict the high and low amplitude regions respectively. (C) Motif robustness analysis. Each network is classified into a unique category based upon its core network topology. In the box plot, the central bar indicates the median estimate of relative robustness and the upper and lower quartiles correspond to the top and bottom of the box. The points correspond to outliers. (D,E) The ten most robust three-node core topologies together with their frequency-amplitude properties.

We find that the ring oscillator core architecture is the most robust, which is a form of delayed negative feedback oscillator. This is followed by topology N2 that can be referred to as an incoherently amplified negative-feedback loop (IANF),^25^ and then N3, which is a combination of a delayed negative feedback (ring) oscillator and two amplified negative feedback oscillators. The N10 system can be considered as as a ring oscillator with additional delayed negative feedback; a design principle that has been shown to be at the core of the mammalian circadian oscillator.^44,45^ More generally, we observe that the motif of mutual activation and repression *X → Y* ⊣ *X* is an important feature of these high frequency oscillators.

Upon examination of the number of regulatory interactions within each category (Supplementary Figure 10) we can see that the top network, N1, contains the least number of edges (or low complexity). This topology scores highly because our definition of robustness automatically takes into account complexity, and essentially scores the robustness per biochemical reaction. In contrast, networks N8–10, contain a maximal six interactions in their core, plus contain further auto-regulatory loops taking the total number of interactions to nine, indicating a fully connected network. These are penalised for containing a high number of interactions. Since our categories represent averages over the auto-regulatory interactions we examined how negative, positive and mixed auto-regulation featured within a topology by counting the number of motifs where these occurred (Supplementary Figure 10). We found that practically every topology contains some form of auto-regulation. In particular, positive auto-regulation dominates in N1 and N2 as expected. However, N3–7 contain varying amounts of mixed auto-regulatory interactions in addition to pure positive auto-regulation, while in N8–10, mixed and pure negative auto-regulation dominate. These differences between categories are expected since some core topologies, for example N4, are not expected to oscillate without any additional auto-regulatory interactions.

To investigate how the frequency and amplitude properties of the robust oscillators depend on the core architecture we re-sampled the posteriors for the top ten core topologies and re-simulated the dynamics (Figure 5E). We found high reproducibility in the stochastic dynamics (Supplementary Figure 11). No correlations between amplitude and frequency were apparent, indicating there are no explicit tradeoffs within these categories. There appears to be a natural frequency range with these parameter priors that gives oscillations with a time period of between 5 and 20 minutes, and an amplitude range spanning two orders of magnitude. Interestingly, although we required an average amplitude greater than 100 protein molecules, most oscillators have an amplitude much higher than this. This implies that the frequency and amplitude are related, and that requiring a specific frequency implies a specific amplitude range. Here the oscillators are constrained by their frequency properties, which gives rise to oscillators with amplitude properties that easily satisfy our constraints. In topologies N3-5, and to a lesser extent N2,6 and 7, we did observe high variability in the frequency, which implies a high frequency tunability. This is obtained through the addition of positive feedback within the core topology, which effectively speeds up the amplified negative feedback. We investigated this phenomena further by performing a regression of all the parameters on oscillator frequency and found that the most significant parameters were the degradation rates for the mRNA and protein species of B and C, plus the edges *I*_7_ and *I*_9_ (Supplementary Figure 12). The highest frequency oscillators are formed from a combination of high degradation rates and positive (activating) *I*_7_ and *I*_9_

### Both the robustness and the phenotype of the oscillators depend on degradation rates

Given the strong dependence of frequency on degradation rates we decided to investigate how changing the prior assumptions on degradation rates affected our analysis of the three-node categories. We changed the prior so that protein and mRNA degradation rates were reduced by an order of magnitude with prior distribution of *U*(0,1) min^−1^. Figure 6A,B shows the distribution of relative robustness and the most robust 10 network topologies. Interestingly, the ring oscillator is much less robust under this scenario and is ranked 21st, which verifies the observation that this topology requires very high degradation rates to function robustly. We see a corresponding increase in positive feedback within the core topologies which can counteract the smaller degradation rates.

**Figure 6:**
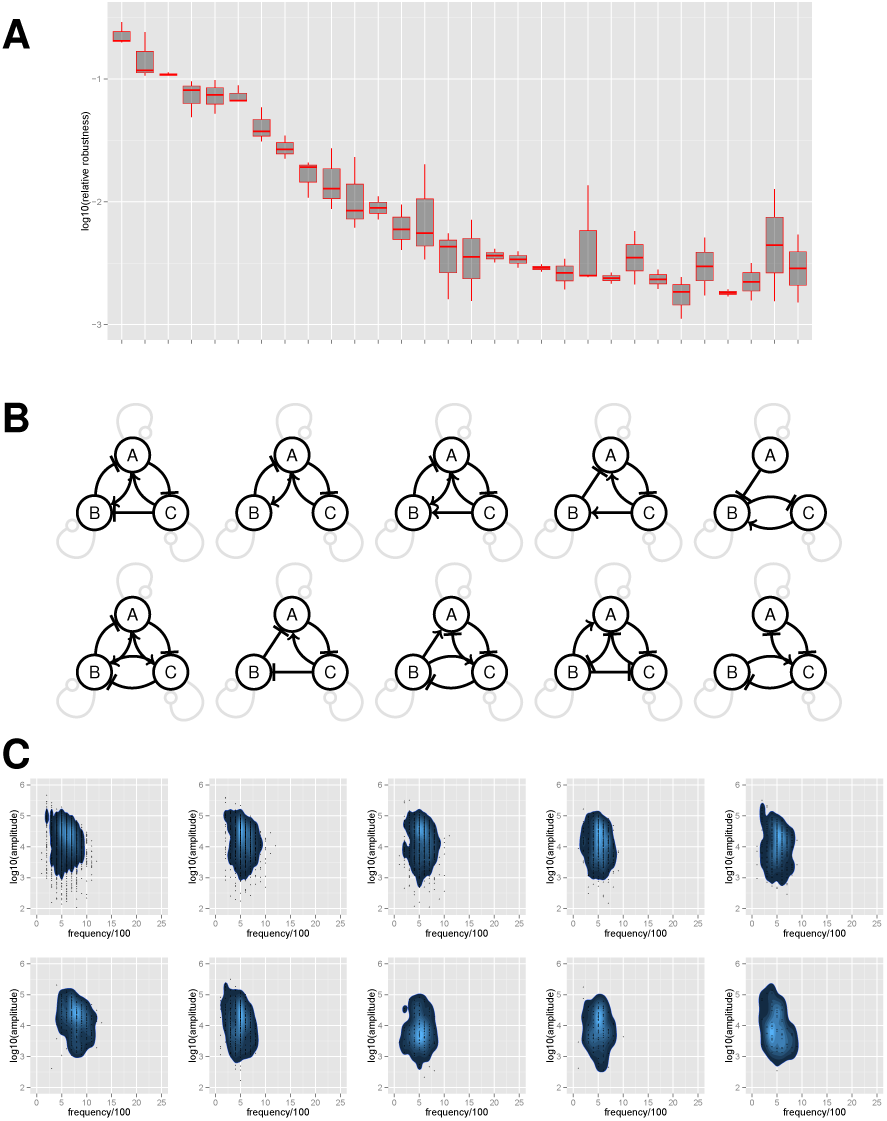
Low frequency oscillators; reanalysis of the three-node networks with a reduced prior on the decay rates of the mRNA and protein species. (A) The distribution of the 30 most robust topologies. In the box plot, the central bar indicates the median estimate and the upper and lower quartiles correspond to the top and bottom of the box. The points correspond to outliers. (B,C) The ten most robust three-node core topologies together with their frequency-amplitude properties.

The frequency and amplitude properties of these oscillators are shown in Figure 6C. We see a doubling in oscillator time period, accompanied by an increase in species number, albeit with a similar amplitude range of two orders of magnitude (see Supplementary Figure 13). We also note the oscillators that previously showed a large variation in frequency no longer demonstrate this behaviour. The frequency and amplitude properties of four different oscillator systems (two-node, ring, three-node and four node) show no significant differences in oscillator frequency and amplitude (see Supplementary Figure 13). These findings suggest that degradation rates are key in defining the frequency and amplitude properties of general oscillators and also that reducing frequency generally implies an increase in amplitude.

That robustness and phenotype of oscillators are affected by degradation rates should come as no surprise. Wong *et. al.* showed that the robustness of their gene-metabolic oscillator depended on the nature of protein degradation^46^ and Stricker *et. al.* added *ssrA* tags to construct their robust two-gene oscillator.^17^ What we have shown is that this dependence applies to a large class of transcriptional oscillators. It also demonstrates the importance of engineering protein degradation in real systems,^47^ which could be applied to the construction of robust oscillators and synthetic circuits more generally.

### Increasing the number of regulatory interactions increases robustness

While it is known from quantitative biology that feedback interactions can increase robustness,^8,9,35,38^ these studies have mostly focussed on a small number of regulatory interactions. Here we examined directly how the robustness of the oscillator systems change as the number of regulatory interactions increases (Figure 7). To do this we examined oscillators formed from a four node interconnected network with 76 parameters (Figure 7A). Again we precomputed the prior on the model space by removing mirror images (transforming only one node) and removing unconnected networks (topologies where the output node was unconnected), which resulted in 7559460 networks (See Supplementary Information). We calculated the relative robustness by taking the ratio of the model posterior probability and the induced prior due to the number of topologies with the given number of interactions (Supplementary Figure 14).

**Figure 7:**
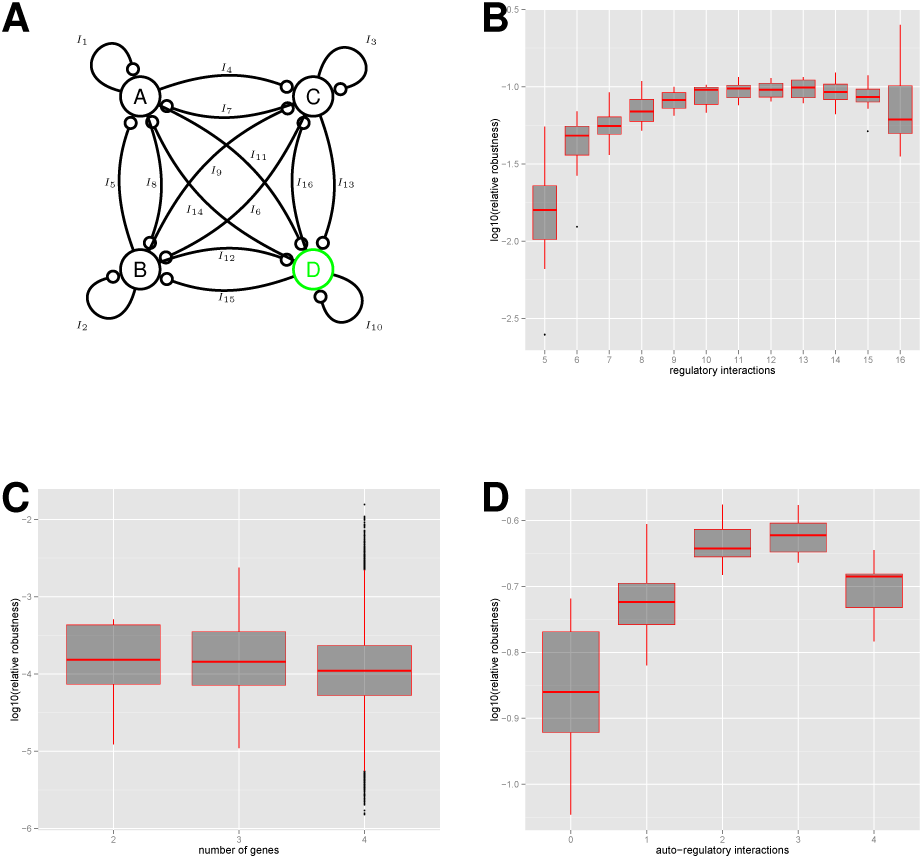
Analysis of robustness and system topology. (A) The regulatory interactions considered for the four-node oscillator system. Oscillator robustness as a function of (B) total number of regulatory interactions, (C) the number of genes and (D) the number of auto-regulatory interactions. In the box plots, the central bar, top and bottom of the box indicates the median, upper and lower quartiles respectively. The points correspond to outliers.

We find that robustness increases with the number of regulatory interactions but only up to around 10 or 11 where it reaches a plateau (Figure 7B). The Bayes factor between 5 interactions and 11 interactions is around 6.5 indicating a substantial increase in robustness. After the plateau is reached, any increase in robustness by adding regulatory interactions approximately equals the increase in the (multi dimensional) volume of parameter space. Another way to express this is that the robustness per biochemical parameter is constant. The implications of this are that while increasing the number of interactions above 11 will possibly produce systems that have an increased level of robustness, the increase in robustness will be directly proportional to the complexity (number of interactions). If we assume that increased complexity comes at a high price (as is generally the case in synthetic biology) then we may conclude that systems with 10 or 11 interactions provides a best case scenario (when constrained to a maximum of four genes). We may also speculate that a similar tradeoff could apply to naturally evolved systems and could be investigated further. Interestingly we don’t see a strong dependence of robustness on the number of genes in the system (Figure 5C). We also split the regulatory interactions into those that comprise the core topology and those that comprise auto-regulatory interactions (Supplementary Figure 14, Figure 5D). We see a similar plateau in the number of core regulatory interactions but see a drop in robustness in the four node system with four auto-regulatory interaction, although the Bayes factor indicates this is not a strong effect. Although we can conclude that increasing the number of genes from two to four does not seem to give any increase in robustness, we cannot rule out that systems with five or more genes show a stronger increase in robustness, and also that the relationship between robustness and regulatory interactions is different in that case.

## Conclusion

We have presented a mathematical framework for the modelling and analysis of robustness in stochastic biological systems and applied it to the case of stochastic transcriptional oscillators. This framework can be used for the design of systems for synthetic biology in order to predict the most robust systems to construct. It can also be used to gain understanding of natural biological systems. Our analysis of transcriptional oscillators has provided a number of insights. We have shown that the oscillator constructed by Stricker *et. al*.^17^ is the most robust two gene system and more robust than the ring oscillator, a fact alluded to but never directly shown.^20^ In addition, we verified the increase in robustness of ring oscillators by the addition of positive feedback,^35^ but also show how this can be achieved in a number of ways and how each of these affects robustness. We searched model space for the most robust three gene systems possible and arrived at systems comprising known and novel oscillator topologies. Our results also indicate that once the structure of the system is fixed, it is the degradation of the species, rather than than the regulatory parameters, that determine oscillator phenotype and robustness. Finally we show that increasing the number of regulatory interactions increases the robustness of the oscillator systems up to a certain level of complexity (number of regulatory interactions). This provides evidence that natural systems should display high levels of interconnectedness to increase robustness, but also that there is a natural limit, beyond which adding further feedback makes no difference. This also suggests that the future of synthetic gene networks lies in the creation of systems comprising additional connectivity to ensure their function across a wide range of conditions. We have highlighted interesting network topologies for further study in order to gain a deeper understanding of their dynamics (stochastic behaviour, robustness, chaos^25^), but also made quantitative predictions on the robustness of different designs, which will be tested in real systems in future work. Realising these oscillating systems and testing the predictions of the framework will further our knowledge of the intricate details of biological systems and allow us to generate accurate but efficient model descriptions.

We believe that our approach has wide applicability across quantitative and synthetic biology, however there are a number of limitations. Our conclusions depend on the modelling assumptions, and here we have tried to find a balance between simplicity and explanatory power with as much biological detail as possible. We did not explicitly include time delays in our model although it is known to be important for oscillating systems.^17,45^ Despite this, the model recapitulates known results, suggesting that our model of mRNA dynamics with large parameter priors causes sufficient delay in the system, although this is not expected to be generally valid^25^ and is also dependent other parameter values.^31^ A related issue is the prior parameter distributions which we generally treat as uniform over a biologically plausible range. As our quantitative knowledge improves, for example with high throughput measurement of biochemical parameters,^48^ this information can be incorporated into our prior, which should be seen as an advantage of the Bayesian approach. The algorithm is computationally intensive and requires high performance computing to explore the kinds of model spaces that are of interest. While the SMC algorithm has desirable properties regarding convergence to the true target distribution in the limit of number of particles, there is still the possibility of falling into local minima. This can be alleviated by running multiple replicates and increasing the number of particles, as in this study, but there is always the possibility that some part of the space was not explored. Finally it is worth emphasising that our framework is probabilistic in nature and its predictions can be interpreted as an average over implementations (specific promoters, genes). Any particular implementation of a single system may possess properties that make it more or less robust than predicted (for example due to temperature dependence^49^).

Future developments of our framework will be to incorporate contextual effects - known to strongly influence gene network dynamics - such as coupling with growth dynamics^50^ and competition for resources.^51–53^ Eventually, this type of approach could be combined with whole cell models.^54,55^ The framework will also be used to investigate the design principles of more general pulsatile systems,^56,57^ investigate systems that provide robust sensing, and also devices for the robust delivery of therapeutic molecules.

Predictive mathematical modelling is at the core of engineering principles and forward design. Currently we lack the tools and knowledge to achieve this in biological systems. Our statistical framework is a significant step in this direction and can provide an engineer with the most robust systems achieving a particular design objective, together with novel and testable predictions. We believe that due to the uncertainty inherent in biological systems, which arises from their complexity and stochasticity, coupled with our lack of detailed knowledge of both their structure and underlying parameters, probabilistic modelling will have a major part to play in the future engineering of biological systems.

## Methods

### Defining and calculating robustness

Intuitively we can think of robustness as a measure of the average performance of a system over some space of perturbations.^6^ Generally speaking, perturbations can be applied at the mutational level, which change the structure of the system, or at the parametric level through changes in rates due to internal and environmental interactions. Consider a model of a system that is required to perform some objective behaviour, *O*. We define an evaluation function or measure of performance, *D*(*x,O*), under the effect of a perturbation *x ∈ χ* with probability distribution given by *p*(*x*). We can then define a general form for the robustness

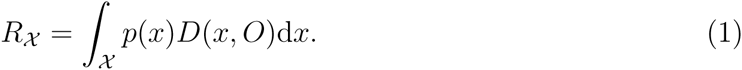

In the case of systems and synthetic biology the model often takes the form of a biochemical reaction system, *M*, with an associated set of reaction rate constants *θ ∈* Θ*_M_*. Here we will focus on robustness to changes in the biochemical parameters and, as in previous studies, will assume that changes to the parameters, *θ*, occur through processes such as the changing of cell size, temperature variations and cellular contexts.^58^ Once this mapping has been applied, the probability distribution of perturbations, *p*(*x*) becomes the probability distribution of reasonable, or physically constrained, parameter values over the range of fluctuations and contexts, *p*(*θ|M*), which in Bayesian statistics is known as the prior distribution (note we have kept the explicit dependence on *M*). In principle we are free to choose any evaluation function for *D*(*θ, O*), however a particularly useful choice is the probability of observing the objective behaviour given the value of *θ*, denoted *p*(*O|θ, M*). Taken as a function of *θ* this is known as the likelihood. The definition of robustness can then be expressed as

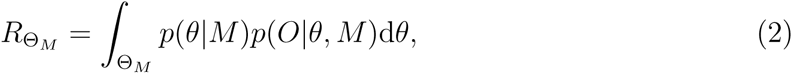

which is precisely the model evidence (or marginal likelihood) from Bayesian statistics, *p*(*O|M*). Note this quantity is also the normalisation constant of the *posterior distribution, p*(*θ|O, M*), which is the probability distribution of the parameters that give rise to the objective behaviour, *p*(*θ|O, M*) ∝ *p*(*θ*)*p*(*O|θ, M*). It is this adoption of the likelihood as our evaluation function that allows us to interpret our results in a probabilistic manner and also provides us with established methods to calculate both *p*(*θ*|*O, M*) and *R*. One advantage in defining *R* as the integral over Θ is that increasing the size of a model (also known as its complexity) increases the size of Θ. This automatically penalises model complexity unless there is a corresponding increase in performance (likelihood) (see Supplementary Information).

For a typical biochemical system of interest Θ is high dimensional. This makes analytical calculation of Equation 2 impossible for all but the simplest systems. In more realistic examples one must use Monte Carlo methods such as Markov chain Monte Carlo (MCMC) or sequential Monte Carlo (SMC) to either calculate *R* directly, or indirectly through first calculating the posterior *p*(*θ*|*O,M*). A further complication arises when the biochemical system is stochastic and represented by either a continuous time Markov process or a system of stochastic differential equations (SDEs). In these cases, *p*(*O|θ, M*) generally cannot be written in closed form and we must evaluate *R* using approximate methods, known collectively as approximate Bayesian computation (ABC), which use simulations to match model output to data.^59–62^

The Bayesian framework allows us to calculate a numerical value for the difference in robustness between two systems, which is expressed through the Bayes factor.^63^ The Bayes factor for how well two models *M*_1_ and *M*_2_ can achieve an objective behaviour is given by

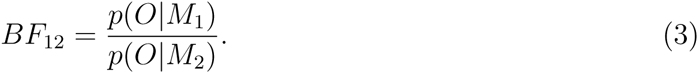

Generally speaking, the value of *BF*_12_ provides evidence that *M*_1_ satisfies the objective behaviour better than *M*_2_, across the parameter space defined by the prior distribution. A *BF*_12_ < 3 indicates weak evidence, *BF*_12_ ≈ 3 − 10 indicates substantial evidence and a *BF* > 10 indicates strong evidence.^63^

Finally it is worth noting that in principle one could use a frequentist model selection framework to calculate robustness, such as the Akaike Information Criterion (AIC).^64^ However, it has been shown previously that this method doesn’t always perform well when applied to the case of discriminating network motif models.^65^

### Representation of network topologies

A given network topology is modelled by a set of reactions describing the interactions of the different mRNA and protein species. Each node in the network represents transcriptional and translational processes, and each edge represents a regulatory interaction. While we expect the conclusions about the robustness of networks to depend strongly on the specific modelling assumptions, in order to create a predictive framework, we must make some modelling decisions that represent some generalities of synthetic gene networks. In the following, we assume that the biochemical system is spatially homogenous, that transcription rates follow Hill-type functions with regulatory proteins acting as dimers, and that protein and mRNA species degrade through first order processes. In reality, the kinetics can be more complex and can include time delays and Michaelis-Menten kinetics for active enzymatic degradation, though in principle these could also be included.

From this set of reactions, and assuming homogenous spatial conditions, a Markov jump process (MJP) governed by the master equation can be formed, which models the probability of each species in the system as a function of time.^66,67^ In principle, this is the most appropriate formalism since it correctly accounts for the probabilistic nature of the dynamics. However, there exists an approximation to the master equation, know as the diffusion approximation, which allows the derivation of stochastic differential equations (SDE), also know as the chemical Langevin equation.^67,68^ We chose to use this formalism, since their simulation is more amenable to parallelisation on Graphics Processing Units and is therefore much more efficient when simulating ensembles of systems with different parameters.^69^ Although one assumption underlying the SDE representation is a large system size (number of molecules), we found a high correlation between summaries generated from SDEs and the MJP under the same parameters (Supplementary Figure 15). Since we are working in a stochastic modelling setting we use the convention that species amount is measured in numbers of molecules rather than concentrations.

Given a node X, we represent the numbers of mRNA and protein molecules as *X_m_RNA* and *X* respectively. Both species have production and decay terms, giving rise to the following SDEs

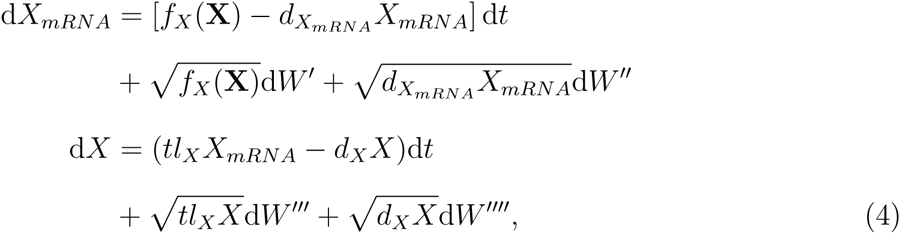

where 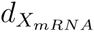, *tl_X_*, *d_X_* are the rate constants for mRNA degradation, translation and protein degradation respectively (all units of min^−1^), and the W represent the stochastic nature of the reactions (independent Wiener processes). Therefore the model of a n node network consists of a system of 2*n* SDEs.

In Equation 4 the regulatory interactions are encoded in the function *f_X_* (X), which represents the binding of proteins to the promoter region upstream of the coding region of X. Here **X** represents all the proteins in the network so, for example, in a two-node network with nodes A and B, *X* = {*A, B*}. To model the probability that a promoter is in the open configuration we utilise the Shea-Ackers formalism, which adopts a thermodynamic argument under equilibrium.^70^ We assume that the regulatory proteins form dimers, which coarsely models commonly used promoter interactions in existing prokaryotic synthetic gene networks, and is important for capturing the nonlinearity present in genetic systems.^17,71^ We further assume that the dimerisation process is in equilibrium

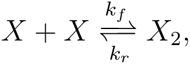

which allows the identification *X*_2_ = *X*^2^*/K_d_* where *K_d_ = k_r_/k_f_* N is the dissociation constant (where the notation, N, corresponds to the integer number of molecules). As an example, given two edges *A → B* and *C* ⊣ *B*, the expression for *f_B_*(*A, B,C*) is given by

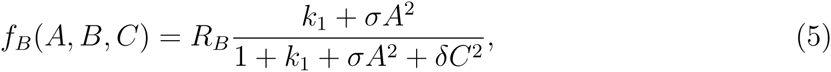

where *k*_1_*,σ* and *δ* are parameters for the relative affinity of RNAP (baseline expression, dimensionless), the affinity of activator *A*_2_ in units N^−2^ and the affinity of repressor *C*_2_ in units N^−2^ respectively (see Supplementary Information). The strength of the promoter is represented by R*_B_* and is measured in units of N min^−1^. The advantages of this modelling formalism is that multiple protein bindings can be handled in a straightforward manner and these parameters are often measured when promoters are characterised, and are therefore available in the literature.^72^

The prior parameter distributions are specified through uniform distributions to reflect the wide range of possible biochemical parameters. Translation and decay rates are given prior distributions of *U*(0,10) min^−1^. The promoter parameters for transcription factor binding (corresponding to *σ* and *δ* in Equation 5) are given priors of *U*(0,1 × 10^6^) N^−2^. The parameters representing baseline expression (corresponding to *k*_1_ in Equation 5) are given priors of *U*(0, 500). Promoter strengths are given priors of *U*(0, 10000) N min^−1^ to reflect variations seen in real synthetic promoters.^72^ The initial conditions of the network must also be specified, which themselves can be treated as parameters. We assign a prior of *U*(0, 100) N on the initial conditions of all mRNA and protein molecules.

### Defining the objective for stochastic oscillations

Oscillatory behaviour in deterministic dynamical systems can be specified in various ways including objective functions minimising squared deviations at regular intervals, Fourier spectra and through the eigenvalues of the Jacobian matrix of the linearised system near the (unstable) steady state.^28,40,41,73^ Quantifying stochastic oscillations presents a challenge since in general the behaviour can range from periodic with constant amplitude to pulsatile with varying time period and amplitude, depending on the level and nature of the noise.^17^

Here we take a signal processing approach to identifying stochastic oscillations. All systems were simulated for *t* = 100 minutes. The raw signal for the protein of interest is smoothed by applying to the time series a mean filter via convolution (Figure 1C). The signal is then differentiated and maxima and minima identified above a signal-to-noise threshold. The locations of these stationary points are used to define the summary statistics *n_max_, n_min_,* which are the number of maxima and minima respectively, and *x_max_,_i_, x_min,k_*, which are the signal level at maxima *i* and minima *k* respectively. The objective behaviour is then specified via distances defined on the summary statistics. For example, in the case where a fixed number of regular pulses is desired, we use the following vector valued distance

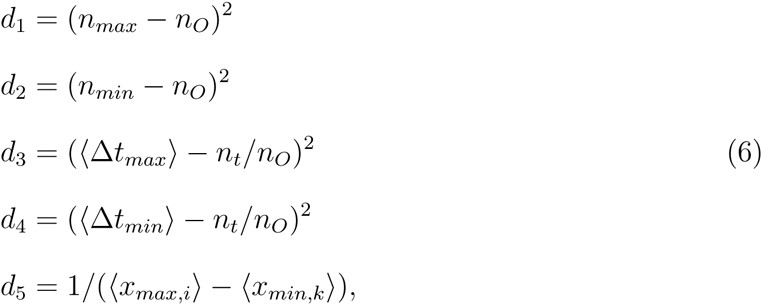

where *n_O_* is the number of desired pulses (here we set *n_O_* = 10), 〈Δ*t_max_*〉, 〈Δ*t_min_*〉 are the mean number of time points between the maxima and minima respectively, *n_t_* is the total number of time points and 〈*x_max_,_i_*〉, 〈*x_min_,_k_*〉 are the average amplitude of the maxima and minima respectively. The distances *d*_1_ and *d*_2_ quantify how close the observed number of peaks are to the objective, *d*_3_ and *d*_4_ quantify the average distance between pulses, and *d*_5_ quantifies the amplitude. The algorithm minimises all distances simultaneously. In this work we assume that the objective is reached when **d** < *∊* where *∊* = (0, 1, 0.1, 0.1, 0.01). This gives rise to regular oscillations with an amplitude of > 100 molecules and a frequency of 0.1 oscillations per minute. We refer to this objective as S1.

In addition to the objective behaviour defining a fixed number of pulses, as above, we also developed an alternative objective defining regular oscillations. In this case the number of desired pulses is free to change

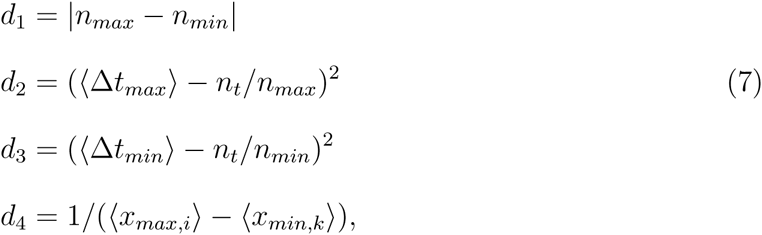

where the main difference is that the number of maxima and minima are constrained to be equal through *d*_1_. In this case, the objective is reached when *∊* = (1, 0.1, 0.1, 0.01) giving rise to regular oscillations with an amplitude of > 100 and a variable frequency. We refer to this objective as S2.

### Maximising robustness simultaneously in model-parameter space

To search jointly through topologies and parameters that satisfy the objective behaviour we developed an extension to the ABC SMC algorithm,^59,60^ inspired by variable selection in a linear regression setting, and implemented within the existing tool ABC-SysBio^61,62^ The total space of possible models is specified through a connected network that contains within it all possible models (Figure 1B). Each edge has an associated integer valued parameter, *I_j_ ∈* {−1, 0, +1}, representing a repressing, missing and activating regulation respectively. The set of *I_j_* are treated as parameters to be inferred from the objective behaviour in an analogous manner to the biochemical rate constants. Our approach differs from other implementations of ABC for network inference,^74^ because we approximate the joint model parameter space as a product, *p*(*M, θ*) = *p*(*M*)*p*(*θ|M*) *≈ p*(*θ*)*p*(*M*). (See Supplementary Information).

The algorithm proceeds through a series of intermediate distributions defined by a decreasing distance to the objective behaviour. Each new weighted population of parameters is obtained from the previous one by resampling, perturbing and performing ABC with a new distance threshold calculated from the previous distribution of distances. Biochemical parameters are perturbed using a uniform distribution while edges in the network are perturbed using a categorical distribution. It is precisely this structure that allows the exploration of model and parameter space simultaneously. The algorithm terminates when the desired distance threshold, *∊*, is reached. The relative evidence of each model present within the posterior distribution is calculated by summing all the corresponding weights. Because the algorithm explores models in direct proportion to their robustness, it focusses in on systems where robustness is high and ignores models where the robustness is negligible.

The Bayesian framework allows us to place priors on model space; we can set the priors for particular models to be zero and therefore reduce the size of the search space. For example, the space of all possible three-node networks contains 3^9^ = 19683 models. However if the output is fixed on one node, say node *C*, then nodes *A* and *B*, are interchangeable. There are 243 networks that are invariant under this symmetry operation with 19440 containing a mirror image, leaving 9963 possible non redundant networks (see Supplementary Information). This situation is handled in a straightforward manner by precomputing the prior over model space. In principle, topologies could also be weighted according to other information rather than simply included and excluded.

## Acknowledgement

C.P.B. and M.L.W. acknowledge funding from the Wellcome Trust through a Research Career Development Fellowship (097319/Z/11/Z). M.L acknowledges funding through the UCL Impact Award scheme. R.P. acknowledges funding from the Wellcome Trust (098325). All the authors would like to acknowledge that the work presented here made use of the Emerald High Performance Computing facility made available by the Centre for Innovation. The Centre is formed by the universities of Oxford, Southampton, Bristol, and University College London in partnership with the STFC Rutherford-Appleton Laboratory.

## Supporting Information Available

The supporting information contains supplementary methods and results. This material is available free of charge via the Internet at http://pubs.acs.org/.

## Graphical TOC Entry

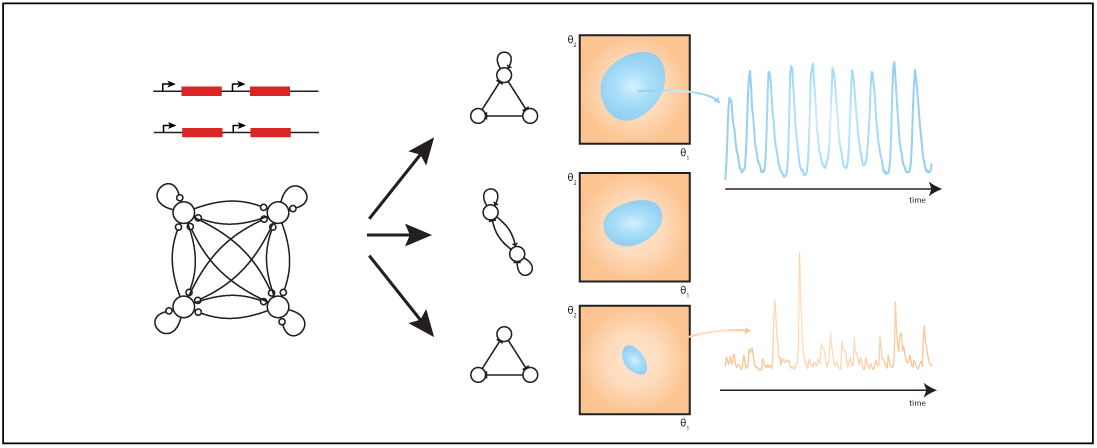

## References

(1) Arkin, A. P. A wise consistency: engineering biology for conformity, reliability, predictability. Current opinion in chemical biology 2013,

(2) Barkai, N.; Leibler, S. Robustness in simple biochemical networks. Nature 1997, 387, 913–917.

(3) Stelling, J.; Sauer, U.; Szallasi, Z.; Doyle, F. J.; Doyle, J. Robustness of cellular functions. Cell 2004, 118, 675–685.

(4) Prill, R. J.; Iglesias, P. A.; Levchenko, A. Dynamic properties of network motifs contribute to biological network organization. PLoS Biology 2005, 3, e343.

(5) Kim, J.; Bates, D.; Postlewaite, I.; Ma, L.; Iglesias, P. Robustness analysis of biochemical network models. Systems Biology 2006, 153, 96.

(6) Kitano, H. Towards a theory of biological robustness. Molecular Systems Biology 2007, 3.

(7) Iglesias, P.; Ingalls, B. Control Theory and Systems Biology; MIT Press, 2010.

(8) Becskei, A.; Serrano, L. Engineering stability in gene networks by autoregulation. Nature 2000, 405, 590–593.

(9) Doyle, J.; Csete, M. Motifs, control, and stability. PLoS Biology 2005, 3, e392.

(10) Kitano, H. Violations of robustness trade-offs. Molecular Systems Biology 2010, 6, 384.

(11) Chandra, F. A.; Buzi, G.; Doyle, J. C. Glycolytic oscillations and limits on robust efficiency. Science 2011, 333, 187–192.

(12) Blanchini, F.; Franco, E. Structurally robust biological networks. BMC systems biology 2011, 5, 74.

(13) Ingram, P. J.; Stumpf, M. P.; Stark, J. Network motifs: structure does not determine function. BMC Genomics 2006, 7, 108.

(14) Elowitz, M. B.; Leibler, S. A synthetic oscillatory network of transcriptional regulators. Nature 2000, 403, 335–338.

(15) Atkinson, M. R.; Savageau, M. A.; Myers, J. T.; Ninfa, A. J. Development of genetic circuitry exhibiting toggle switch or oscillatory behavior in Escherichia coli. Cell 2003, 113, 597–607.

(16) Fung, E.; Wong, W. W.; Suen, J. K.; Bulter, T.; Lee, S.-g.; Liao, J. C. A synthetic gene-metabolic oscillator. Nature 2005, 435, 118–122.

(17) Stricker, J.; Cookson, S.; Bennett, M. R.; Mather, W. H.; Tsimring, L. S.; Hasty, J. A fast, robust and tunable synthetic gene oscillator. Nature 2008, 456, 516–519.

(18) Tigges, M.; Marquez-Lago, T. T.; Stelling, J.; Fussenegger, M. A tunable synthetic mammalian oscillator. Nature 2009, 457, 309–312.

(19) Kim, J.; Winfree, E. Synthetic in vitro transcriptional oscillators. Molecular Systems Biology 2011, 7, 465.

(20) Purcell, O.; Savery, N. J.; Grierson, C. S.; di Bernardo, M. A comparative analysis of synthetic genetic oscillators. Journal of the Royal Society, Interface / the Royal Society 2010,

(21) Danino, T.; Mondragón-Palomino, O.; Tsimring, L.; Hasty, J. A synchronized quorum of genetic clocks. Nature 2010, 463, 326–330.

(22) Mondragón-Palomino, O.; Danino, T.; Selimkhanov, J.; Tsimring, L.; Hasty, J. Entrainment of a population of synthetic genetic oscillators. Science 2011, 333, 1315–1319.

(23) Prindle, A.; Samayoa, P.; Razinkov, I.; Danino, T.; Tsimring, L. S.; Hasty, J. A sensing array of radically coupled genetic ‘biopixels’. Nature 2012, 481, 39–44.

(24) Goldbeter, A. Computational approaches to cellular rhythms. Nature 2002, 420, 238–245.

(25) Novák, B.; Tyson, J. J. Design principles of biochemical oscillators. Nature Reviews Molecular Cell Biology 2008, 9, 981–991.

(26) Lenz, P.; Søgaard-Andersen, L. Temporal and spatial oscillations in bacteria. Nature reviews Microbiology 2011, 9, 565–577.

(27) Vilar, J. M. G.; Kueh, H. Y.; Barkai, N.; Leibler, S. Mechanisms of noise-resistance in genetic oscillators. Proceedings of the National Academy of Sciences of the United States of America 2002, 99, 5988–5992.

(28) François, P.; Hakim, V. Design of genetic networks with specified functions by evolution in silico. Proceedings of the National Academy of Sciences of the United States of America 2004, 101, 580–585.

(29) Guantes, R.; Poyatos, J. F. Dynamical principles of two-component genetic oscillators. PLoS Computational Biology 2006, 2, e30.

(30) Bailey, M.; Joo, J. Identification of network motifs capable of frequency-tunable and robust oscillation. arXiv.org 2011,

(31) Lomnitz, J. G.; Savageau, M. A. Strategy Revealing Phenotypic Differences among Synthetic Oscillator Designs. ACS synthetic biology 2014, 3, 686–701.

(32) Wagner, A. Circuit topology and the evolution of robustness in two-gene circadian oscillators. Proceedings of the National Academy of Sciences of the United States of America 2005, 102, 11775–11780.

(33) Ghaemi, R.; Sun, J.; Iglesias, P. A.; Del Vecchio, D. A method for determining the robustness of bio-molecular oscillator models. BMC systems biology 2009, 3, 95.

(34) Zamora-Sillero, E.; Hafner, M.; Ibig, A.; Stelling, J.; Wagner, A. Efficient characterization of high-dimensional parameter spaces for systems biology. BMC systems biology 2011, 5, 142.

(35) Tsai, T. Y.-C.; Choi, Y. S.; Ma, W.; Pomerening, J. R.; Tang, C.; Ferrell, J. E. Robust, tunable biological oscillations from interlinked positive and negative feedback loops. Science 2008, 321, 126–129.

(36) Nguyen, L. K. Regulation of oscillation dynamics in biochemical systems with dual negative feedback loops. Journal Of The Royal Society Interface 2012, 9, 1998–2010.

(37) Pokhilko, A.; ndez, A. P. n. a. F. a.; Edwards, K. D.; Southern, M. M.; Halliday, K. J.; Millar, A. J. The clock gene circuit in Arabidopsis includes a repressilator with additional feedback loops. Molecular Systems Biology 2012, 8, 1–13.

(38) Teng, S.-W.; Mukherji, S.; Moffitt, J. R.; de Buyl, S.; O’Shea, E. K. Robust circadian oscillations in growing cyanobacteria require transcriptional feedback. Science 2013, 340, 737–740.

(39) Castillo-Hair, S. M.; Villota, E. R.; Coronado, A. M. Design principles for robust oscillatory behavior. Systems and synthetic biology 2015, 9, 125–133.

(40) Barnes, C. P.; Silk, D.; Sheng, X.; Stumpf, M. P. H. Bayesian design of synthetic biological systems. Proceedings of the National Academy of Sciences of the United States of America 2011, 108, 15190–15195.

(41) Barnes, C. P.; Silk, D.; Stumpf, M. P. H. Bayesian design strategies for synthetic biology. Interface focus 2011, 1, 895–908.

(42) Hasty, J.; Dolnik, M.; Rottschäfer, V.; Collins, J. J. Synthetic gene network for entraining and amplifying cellular oscillations. Physical Review Letters 2002, 88, 148101.

(43) Turcotte, M.; Garcia-Ojalvo, J.; Süel, G. M. A genetic timer through noise-induced stabilization of an unstable state. Proceedings of the National Academy of Sciences of the United States of America 2008, 105, 15732–15737.

(44) Hogenesch, J. B.; Ueda, H. R. Understanding systems-level properties: timely stories from the study of clocks. Nature Reviews Genetics 2011, 12, 407–416.

(45) Ukai-Tadenuma, M.; Yamada, R. G.; Xu, H.; Ripperger, J. A.; Liu, A. C.; Ueda, H. R. Delay in feedback repression by cryptochrome 1 is required for circadian clock function. Cell 2011, 144, 268–281.

(46) Wong, W. W.; Tsai, T. Y.; Liao, J. C. Single-cell zeroth-order protein degradation enhances the robustness of synthetic oscillator. Molecular Systems Biology 2007, 3, 130.

(47) Cameron, D. E.; Collins, J. J. Tunable protein degradation in bacteria. Nature Biotechnology 2014, 32, 1276–1281.

(48) Geertz, M.; Shore, D.; Maerkl, S. J. Massively parallel measurements of molecular interaction kinetics on a microfluidic platform. Proceedings of the National …. 2012.

(49) Chandraseelan, J. G.; Oliveira, S. M. D.; Häkkinen, A.; Tran, H.; Potapov, I.; Sala, A.; Kandhavelu, M.; Ribeiro, A. S. Effects of temperature on the dynamics of the LacI-TetR-CI repressilator. Molecular BioSystems 2013, 9, 3117–3123.

(50) Klumpp, S.; Zhang, Z.; Hwa, T. Growth Rate-Dependent Global Effects on Gene Expression in Bacteria. Cell 2009,

(51) Cookson, N. A.; Mather, W. H.; Danino, T.; Mondragón-Palomino, O.; Williams, R. J.; Tsimring, L. S.; Hasty, J. Queueing up for enzymatic processing: correlated signaling through coupled degradation. Molecular Systems Biology 2011, 7, 561.

(52) Prindle, A.; Selimkhanov, J.; Li, H.; Razinkov, I.; Tsimring, L. S.; Hasty, J. Rapid and tunable post-translational coupling of genetic circuits. Nature 2014, 508, 387–391.

(53) Weisse, A. Y.; Oyarzún, D. A.; Danos, V.; Swain, P. S. Mechanistic links between cellular trade-offs, gene expression, and growth. Proceedings of the … 2015,

(54) Karr, J. R.; Sanghvi, J. C.; Macklin, D. N.; Gutschow, M. V.; Jacobs, J. M.; Bolival, B.; Assad-Garcia, N.; Glass, J. I.; Covert, M. W. A whole-cell computational model predicts phenotype from genotype. Cell 2012, 150, 389–401.

(55) Purcell, O.; Jain, B.; Karr, J. R.; Covert, M. W.; Lu, T. K. Towards a whole-cell modeling approach for synthetic biology. Chaos (Woodbury, NY) 2013, 23, 025112.

(56) Locke, J. C. W.; Young, J. W.; Fontes, M.; Jimenez, M. J. H.; Elowitz, M. B. Stochastic Pulse Regulation in Bacterial Stress Response. Science 2011, 334, 366–369.

(57) Levine, J. H.; Lin, Y.; Elowitz, M. B. Functional roles of pulsing in genetic circuits. Science 2013, 342, 1193–1200.

(58) Toni, T.; Tidor, B. Combined model of intrinsic and extrinsic variability for computational network design with application to synthetic biology. PLoS Computational Biology 2013, 9, e1002960.

(59) Toni, T. Approximate Bayesian computation for parameter inference and model selection in systems biology. Thesis 2010, 1–168.

(60) Toni, T.; Stumpf, M. P. H. Simulation-based model selection for dynamical systems in systems and population biology. Bioinformatics 2010, 26, 104–110.

(61) Liepe, J.; Barnes, C.; Cule, E.; Erguler, K.; Kirk, P.; Toni, T.; Stumpf, M. P. H. ABC-SysBio-approximate Bayesian computation in Python with GPU support. Bioinformatics 2010, 26, 1797–1799.

(62) Liepe, J.; Kirk, P.; Filippi, S.; Toni, T.; Barnes, C. P.; Stumpf, M. P. H. A framework for parameter estimation and model selection from experimental data in systems biology using approximate Bayesian computation. Nature protocols 2014, 9, 439–456.

(63) Kass, R.; Raftery, A. Bayes Factors. Journal Of The American Statistical Association 1995, 90, 773–795.

(64) Burnham, K.; Anderson, D. Model Selection and Multimodel Inference: A Practical Information-Theoretic Approach; Springer New York, 2010.

(65) Domedel-Puig, N.; Pournara, I.; Wernisch, L. Statistical model comparison applied to common network motifs. BMC systems biology 2010, 4, 18.

(66) Van Kampen, N. Stochastic Processes in Physics and Chemistry; North-Holland Personal Library; Elsevier Science, 2011.

(67) Wilkinson, D. Stochastic Modelling for Systems Biology, Second Edition; Chapman & Hall/CRC Mathematical and Computational Biology; Taylor & Francis, 2011.

(68) Gillespie, D. T. The chemical Langevin equation. Journal Of Chemical Physics 2000, 113, 297.

(69) Zhou, Y.; Liepe, J.; Sheng, X.; Stumpf, M. P. H.; Barnes, C. GPU accelerated biochemical network simulation. Bioinformatics 2011, 27, 874–876.

(70) Ackers, G. K.; Johnson, A. D.; Shea, M. A. Quantitative model for gene regulation by lambda phage repressor. Proceedings of the National Academy of Sciences of the United States of America 1982, 79, 1129–1133.

(71) Gardner, T. S.; Cantor, C. R.; Collins, J. J. Construction of a genetic toggle switch in Escherichia coli. Nature 2000, 403, 339–342.

(72) Tamsir, A.; Tabor, J. J.; Voigt, C. A. Robust multicellular computing using genetically encoded NOR gates and chemical ‘wires’. Nature 2010,

(73) Chickarmane, V.; Paladugu, S. R.; Bergmann, F.; Sauro, H. M. Bifurcation discovery tool. Bioinformatics 2005, 21, 3688–3690.

(74) Rau, A.; Jaffrézic, F.; Foulley, J.; Doerge, R. Reverse engineering gene regulatory networks using approximate Bayesian computation. Statistics and Computing 2011,

